# Reduced mammillary body volume in individuals with a schizophrenia diagnosis: an analysis of the COBRE data set

**DOI:** 10.1101/2023.06.13.544746

**Authors:** Michal M. Milczarek, Syed I. A. Gilani, Maarten H. Lequin, Seralynne D. Vann

## Abstract

While the frontal cortices and medial temporal lobe are well-associated with schizophrenia, the involvement of wider limbic areas is less clear. The mammillary bodies are important for both complex memory formation and anxiety and are implicated in several neurological disorders that present with memory impairments. However, little is known about their role in schizophrenia. Post-mortem studies have reported both a loss of neurons in the mammillary bodies but also reports of increased mammillary body volume. The findings from *in vivo* MRI studies have also been mixed, but studies have typically only involved small sample sizes. To address this, we acquired mammillary body volumes from the open-source COBRE dataset, where we were able to manually measure the mammillary bodies in 72 individuals with a schizophrenia diagnosis and 74 controls. Participant age ranged from 18-65. We found the mammillary bodies to be smaller in the patient group, across both hemispheres, after accounting for the effects of total brain volume and gender. Hippocampal volumes, but not subiculum or total grey matter volumes, were also significantly lower in patients. Given the importance of the mammillary bodies for both memory and anxiety, this atrophy could contribute to the symptomology in schizophrenia.

## 1. Introduction

Schizophrenia constitutes only a minor proportion of all psychiatric disease cases, however, given the severity and range of associated symptoms, its impact on affected individuals and their families is especially high. Schizophrenia episodes are typically associated with hallucinations and thought disturbances, but individuals are also affected by difficulties in perception, executive function, and other cognitive skills, including memory and emotional processing. While there are treatments for the positive symptoms, there is very little available to alleviate chronic cognitive impairments, which contribute significantly to the long-term impact on patients and their families. Schizophrenia research has focused in large part on the prefrontal cortex, striatum and especially hippocampus (Sigurdsson and Duvarci, 2016). However, it is becoming clear that far more widespread circuits are implicated, contributing to the complexity of the disorder. A better understanding of the circuitry affected in schizophrenia will help improve treatment options.

The mammillary bodies form part of the medial diencephalon and play an important role in the integration of positional, sensory information and arousal (Schaper et al., 2020), which underlies the formation of complex memories. Mammillary body pathology has been reported in multiple neurological disorders (Lequin et al., 2021; Meys et al., 2022; Sheedy et al., 1999), however, the integrity of the mammillary bodies in schizophrenia is not currently clear. Post-mortem studies have assessed the mammillary bodies in individuals with schizophrenia. One study found reduced neuronal density along with enlarged volume in patients with schizophrenia, resulting in no net change in overall neuronal number (Briess et al., 1998). However, data in the patient group in this study exhibited higher variance than in controls and the authors did not provide an age range for the participants. Another post-mortem study found a marked loss of parvalbumin neurons in the mammillary bodies (Bernstein et al., 2007). In both cases, changes were more pronounced in the left hemisphere than the right (Bernstein et al., 2007; Briess et al., 1998). However, since post-mortem studies are often limited to relatively small samples - particularly for the mammillary bodies which are more likely to be damaged on extraction given their location - it has been challenging to relate the observed changes to ageing and other stratifying factors. Furthermore, the mammillary bodies are particularly sensitive to hypoxia, so cell loss may be harder to interpret in post-mortem studies as they can be affected by the cause of death (Kumar et al., 2009; Schmidtke, 1993).

Analysing mammillary body volumes from *in vivo* MRI studies can address some of the limitations with post-mortem studies. One MRI study that assessed mammillary body volume in patients with a schizophrenia diagnosis found a trend for the right mammillary body to be larger in patients with schizophrenia compared to controls (Tognin et al., 2012). A further study used a larger database to assess mammillary body volume in patients with schizophrenia and found no difference (Klomp et al., 2012). However, the inter-rater correlation was quite low for the mammillary bodies, and lower than for other structures in the study, which may have affected the accuracy of mammillary body delineation (Klomp et al., 2012).

As such, based on current studies, the nature of mammillary body pathology in schizophrenia remains unclear. This is likely due to multiple factors, including the heterogeneity of schizophrenia, stratification of the patient population and differences in imaging protocols and/or the segmentation of the mammillary bodies. To address some of these issues, we carried out a volumetric analysis of a large open-source data set (COBRE, see Methods below). Due to the small size of the mammillary bodies and their proximity to the 3^rd^ ventricle (whose size may be affected in schizophrenia), mammillary body volumes were segmented manually by an experienced researcher. Three additional automatically extracted volumes were included in the analyses: the hippocampus, the subiculum, and the total grey matter. The hippocampal formation has been repeatedly shown to be affected in schizophrenia (e.g., Heckers and Konradi, 2002) and it constitutes the primary excitatory input to the mammillary bodies (mainly via the subiculum) (Vann, 2010). Inclusion of total grey matter enabled contrasting regional changes with whole-brain atrophy. To further account for inter-individual differences, we also performed analyses following normalisations to total brain volume and gender, using three different commonly used methods: a ratio method, a covariance method and an allometric method to account for non-linearity in the scaling of sub-cortical regions (Nadig et al., 2018; see Methods for further details). Finally, we also investigated the effect of age on regional volumes.

## 2. Materials and Methods

### COBRE database

The Center for Biomedical Research Excellence (COBRE) contributed a total of 146 structural MR images data from 72 patients with a schizophrenia diagnosis and 74 healthy controls. In the control group there were 23 females (age 18-58) and 51 males (age 18-65; one left-handed). In the patient group there were 14 females (age 20-65; one left-handed) and 58 males (age 18-64; 9 left-handed). Participants had been excluded from the study if they had any neurological disorders, severe head trauma with more than 5 minutes loss of consciousness, history of substance abuse or dependence within the last 12 months. Diagnostic information for this database was collected using the Structured Clinical Interview used for DSM Disorders (SCID).

### MRI acquisition

For the anatomical images, a sagittal five-echo 3D MPRAGE (Multi-Echo MPR) sequence was used with the following parameters: TE1=1.64 ms, TE2=3.5 ms, TE3=5.36 ms, TE4=7.22 ms, TE5=9.08 ms, TR/TI=2530/900 ms, Flip angle=7°, FOV=256×256 mm, Slab thickness=176 mm, Matrix=256×256×176, Series=interleaved, Multislice mode=single shot, Voxel size=1×1×1 mm, Pixel bandwidth=650 Hz. The total scan time was 6 min. TR, TI and time to encode partitions for this acquisition method are similar to that of a conventional MPRAGE, however, with 5 echoes it resulted in visually better grey matter-white matter-cerebrospinal fluid contrast compared to typical MPRAGE.

The imaging data and phenotypic information for COBRE database was collected and shared by the Mind Research Network and the University of New Mexico funded by a National Institute of Health Center of Biomedical Research Excellence (COBRE) grant 1P20RR021938-01A2.

### Mammillary body volumes

All structural 3D volumes were re-sampled on a 0.5 × 0.5 × 0.5 mm grid prior to any volumetric analyses using FSL (FMRIB Software Library) to improve resolution and accuracy. The left and right mammillary bodies were segmented manually using the ITK-SNAP software [version 3.8; www.itksnap.org (Yushkevich et al., 2006)] and volumes were obtained using the same software.

All three planes were used to finalise the segmentations (Figure 1). All segmentations were carried out without knowledge of the demographic data or group allocations. The same experimenter (SDV) carried out all segmentations; repeat segmentations were carried out on a subset of scans to acquire an intra-rater reliability measure. Segmentations were visually inspected by a second experimenter (S.G.) for quality control, again this was done without knowledge of group allocation. Three measures were obtained for each participant: MB_L, MB_R and MB_T corresponding to the left, right and total mammillary bodies.

**Figure 1.**
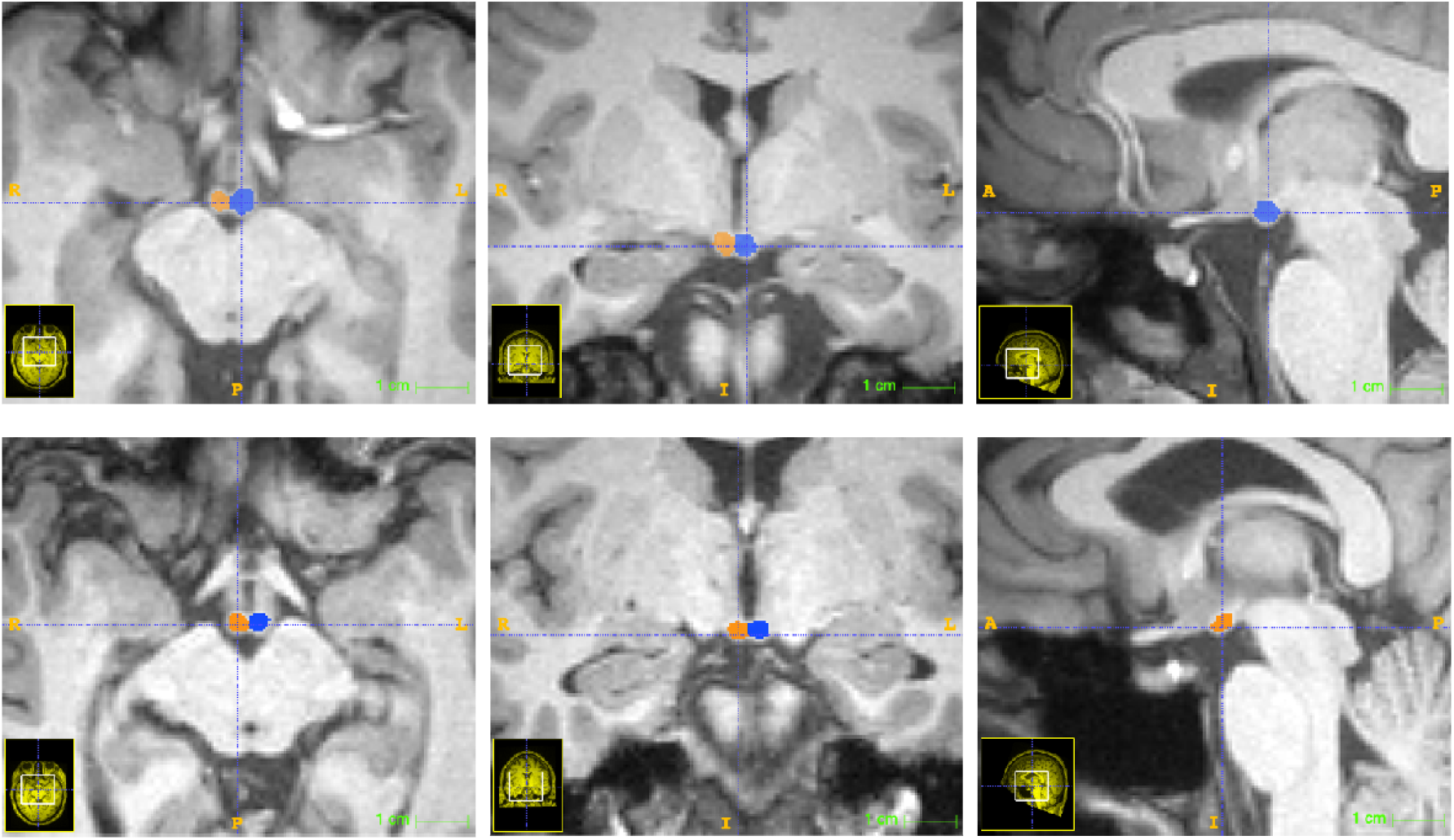
Two examples of mammillary body segmentation using the ITK-SNAP software. Each image row represents axial, coronal and sagittal images from the same brain.

### Whole brain volumes

FreeSurfer (Fischl, 2012) version 7.3 was used with default settings for evaluating total brain volume (Brain Segmentation Volume Without Ventricles) and total grey matter volume using the 3D T_1_-weighted MRI scans.

### Hippocampal volumes

Hippocampal volumes were measured using the FreeSurfer’s subregion segmentation utility provided with FreeSurfer 7.3. The left and right hippocampus volumes were calculated separately for the whole hippocampus, whole hippocampal body, whole hippocampal head, subiculum body and subiculum head subregions. Following extraction of the subregions, they were merged into whole hippocampus (HPC_L, HPC_R) and whole subiculum (SUB_L, SUB_R) for each hemisphere as well as total volumes (HPC_T, SUB_T).

### Statistical analysis

Statistical analysis and data handling were performed in Matlab 2022b (Mathworks, USA) using built-in functions unless otherwise stated.

### Demographic statistics

Ages were compared across groups using a t-test while a Chi-squared test was used to compare gender and handedness distributions across groups.

### Volumetric comparisons

We utilised a generalised linear model (GLM) approach with each subregion modelled separately. Data were analysed for Raw (unadjusted) volumes and following three normalisation procedures to adjust for total brain volume (TBV): the Ratio, Covariance and Allometric techniques (Nadig et al., 2018). The Ratio method involved dividing regional volumes by the TBV for each participant; the Covariance method involved regressing out the effect of TBV so that residuals could be compared and lastly, the Allometric method was analogous to the Covariance method except it used log10 transformed data. The distributions of residuals for each model were visually assessed using Q-Q plots and accepted as approximately normal. The effect of Gender was accounted for where it significantly improved model fits: the most parsimonious model was chosen for each region based on a log likelihood ratio test comparing a model with an interaction term, an additive model and a simple (no Gender as a factor) model. For raw and ratio adjusted volumes, Gender was factored in when testing for the effect of Group whereas for covariance and allometrically adjusted volumes, it was factored in during the normalisation process and then again when testing for the Group effect. The effect of Age was probed in covariance and allometrically adjusted volumes (as Gender was already accounted for in these datasets) following the same iterative model comparison method as described above. Data were also fitted with a segmented regression approach (using the SegReg software; https://www.waterlog.info/segreg.htm), to test whether the rate of volume change differed with age, and split into younger and older cases based on the presence of a significant regression breakpoint.

Since data for some region × condition combinations were not normally distributed, we utilised medians and interquartile ranges as measures of central tendency and variability, respectively. Group differences, where significant, were reported along with median-based effect sizes (Cohen’s d calculated with median and median absolute deviation). All *p*-values were subjected to family-wise multiple comparison correction (Bonferroni method). Data were plotted following z-score transformations (to allow for simultaneous visualisation of vastly different volumes and units, using a median-based, ‘robust’ normalisation method) with gramm (Morel, 2018) in Matlab and modified in PowerPoint (Microsoft Office, USA). An α threshold of 0.05 was selected for significance. All scripts and volumetric data are available upon request.

## 3. Results Demographics

Table 1 displays summary statistics for Age, Gender and Handedness for both groups. The proportion of left-handed individuals was higher in the Patient group (*χ*^*2*^ = 6.3, *p* = 0.012), while Age and Gender did not differ between groups. Since only a very small proportion of individuals displayed left-handedness (<7%), this measure was not included in subsequent analyses.

**Table 1.** Demographic data. Age is mean ± standard deviation. T-values are reported for age and χ^2^ for gender and handedness. L: left handed; R: right-handed; B: use both left and right hands.

| Index | Controls (n = 74) | Patients (n = 72) | t value/$\chi^2$ | p value |
| --- | --- | --- | --- | --- |
| Age | 35.82 ± 11.58 | 38.17 ± 13.89 | -1.11 | 0.27 |
| Gender (F: M) | 23:51 | 14:58 | 2.61 | 0.11 |
| Handedness (L: R: B) | 1:71:2 | 9:61:2 | 6.3 | 0.012 |

### Volumetric differences

We analysed estimated volumes for the mammillary bodies, the hippocampus and the subiculum, as well as total grey matter. All regions other than the mammillary bodies were automatically extracted while the mammillary bodies were manually segmented. The intra-rater reliability measure for manual segmentation was high (0.94).

We utilised unadjusted (Raw) as well as Total Brain Volume-normalised volumes derived using three different methods (Ratio, Covariance, Allometric) to ensure the robustness of the analyses. The effect of Gender was accounted for where it significantly improved the model fit (see Methods). Using Raw volumes, significant differences were found in the mammillary bodies (tleft = -4.25, *p*left = 3.9×10^−4^; tright = -4.7, *p*right = 5.3×10^−5^; ttotal = -4.64, *p*total = 7.6×10^−5^) as well as in the hippocampus (tleft = -1.99, *p*left = 0.01; ttotal = -3.01, *p*total = 0.03), revealing lower volumes in patients with schizophrenia. Normalisation methods preserved the main effect of Group in the mammillary bodies and, for the covariance method, in the hippocampus (see Figure 2 for details). Whereas the hippocampus showed differences only in the left hemisphere (or when analysed as a whole), mammillary body volumes displayed the same degree of volume loss across both hemispheres (allometric method: z = 0.1, *p* > 0.05, z-test) (Paternoster et al., 1998).

**Figure 2.**
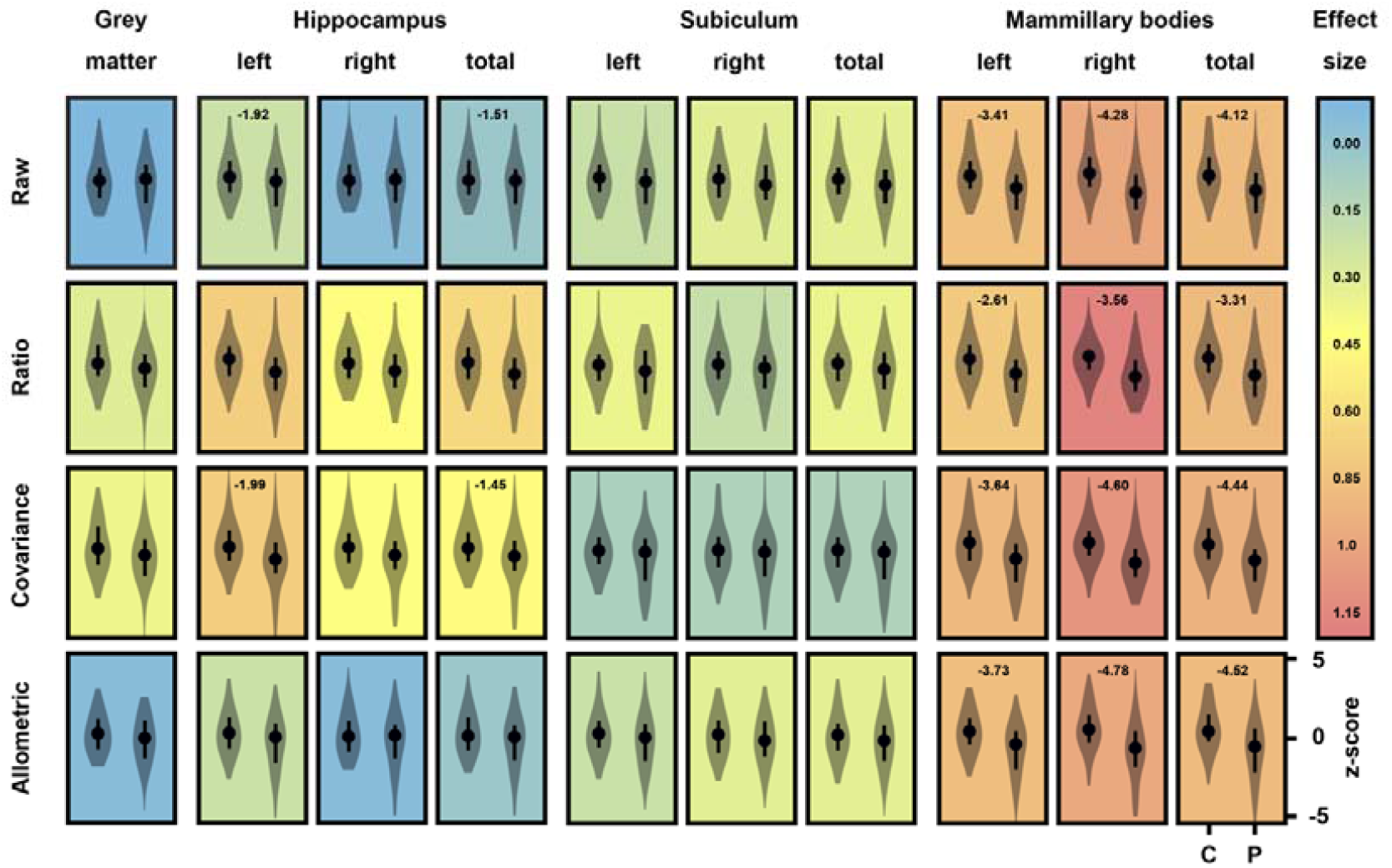
Volumetric differences. The composite chart displays violin plots of z-scored regional brain volumes in Controls (C) and Patients (P) either unadjusted (Raw) or normalised by Total Brain Volume (and Gender) according to the Ratio, Covariance and Allometric techniques (see Methods). The dots represent median values and error bars, the interquartile range. The colour of the background in each plot indicates the effect size of the difference (median-based Cohen’s d) while the numbers on top of the plots are log10 *p*-values for significant differences only.

### Relationship with Age

We further investigated whether the observed volumetric differences in (total) mammillary body volume were age dependent. Based on both covariance and allometrically adjusted volumes, there was a significant Age by Group interaction explaining 16% of the observed variance (t = -2.35/-2.50, *p* = 0.02/0.02 for the covariance and allometric methods, respectively), revealing age-related mammillary body atrophy in Patients with schizophrenia (at a rate of -4.26% per decade). Since the relationship between Age and mammillary body volume in Patients appeared non-linear (Figure 3), we sought to model it with a segmented regression approach (SegReg; https://www.waterlog.info/segreg.htm). Indeed, the rate of volume change differed by Age, as evidenced by the presence of a significant breakpoint (covariance: F(3,67) = 7.6, p < 1.9×10^−4^, allometric: F(3,67) = 5.2, p < 1.9 × 10^−3^). In younger Patients (< 33.04 years old), there was a positive relationship between volume and Age (covariance: t = 4.67, p = 5.49×10_-5_, allometric: t = 3.79, *p* = 6.54×10_-4_) while older Patients (≥ 33.04 years old) did not display any relationship with Age (Figure 2). Control cases did not exhibit a significant breakpoint but a small, continuously positive relationship with Age (covariance method only: t = 2.33, *p* = 0.02). Consistent with this, data split according to the breakpoint revealed volume reductions only in older patients with schizophrenia (t = -5.66/- 5.66, p = 2.56×10^−7^ /2.58×10^−7^ for the covariance and allometric methods, respectively), explaining 29% of the observed variance.

**Figure 3.**
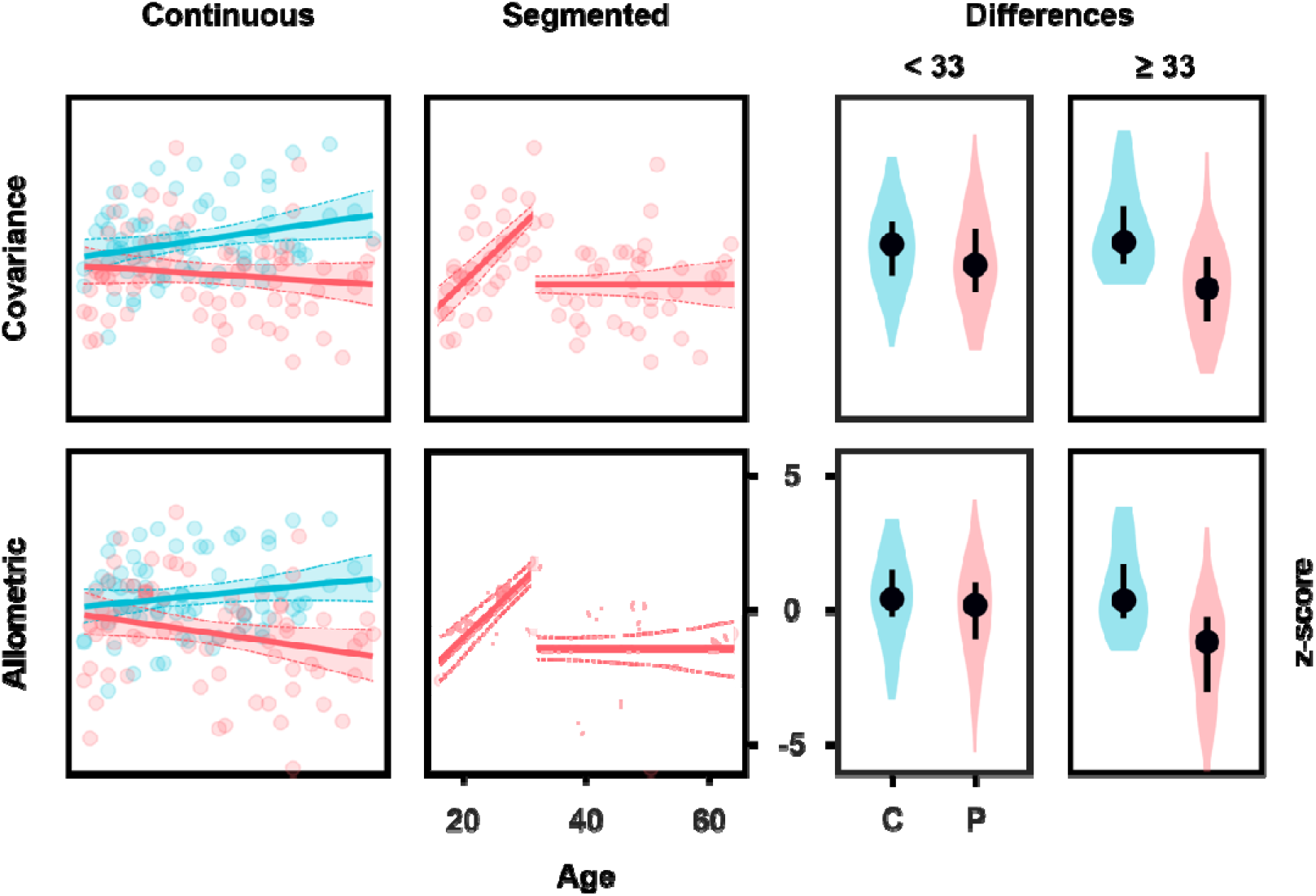
Relationship of mammillary body volume with Age. The scatterplots on the left display fits for covariance and allometric adjusted mammillary body volumes versus Age for continuous regressions for patients and controls and segmented regressions for patients (there was no breakpoint for controls). The shaded areas represent 95% confidence intervals. On the right, the violin plots display data split according to the segmented regression breakpoint at 33 years of age. The dots are medians and error bars, the interquartile range.

## 4. Discussion

Schizophrenia is a complex disorder, and the underlying neural basis is still uncertain. Limbic brain regions have been repeatedly implicated but the findings for the mammillary bodies have been mixed. We made use of a large cross-sectional sample to improve power and to enable us to study the effects of age, gender, and brain hemisphere on mammillary body volume. The results provide evidence for schizophrenia-mediated atrophy of the mammillary bodies (as well as the hippocampus), which is robust against age-related and inter-individual differences in whole-brain volume. While overall grey matter typically decreases at a greater rate in patients with schizophrenia than in controls (Olabi et al., 2011), the mammillary body volume loss we report exceeded that explained by grey matter decline alone.

As with previous studies, the left mammillary body was found to be slightly larger (6.4%) than the right in control participants (Tognin et al., 2012; Vann et al., 2022). In the current data set, the left mammillary body was also larger (6.96%), than the right in the patient group, whereas a previous study reported the left and right mammillary bodies to be of equivalent volumes in patients with schizophrenia (Tognin et al., 2012); a further study did not separate mammillary body volumes by hemisphere (Klomp et al., 2012). While there were hemispheric differences in mammillary body volume in both the patients and controls in the present data set, the hemispheres were not differentially affected in schizophrenia with an equivalent reduction in volume in both the left and the right hemisphere. While other studies have reported increases in volume on either the right (Tognin et al., 2012) or the left (Briess et al., 1998) hemisphere, consistent with the present study, one post-mortem study found bilateral mammillary body neuronal loss, although the loss was larger on the left (Bernstein et al., 2007). A recent study also found a reduction in the dopamine D3 receptor in both left and right mammillary bodies (Mitelman et al., 2020). Together with the present data, this suggests that the mammillary bodies are impacted across both hemispheres in schizophrenia.

There were no significant interactions between mammillary body volume and gender across the patient and control groups. However, numerically, there was a greater volume decrease in male patients (15.5%) compared to female patients (9.5%). As there were more males than females in the present data set, it is possible that with balanced groups, this difference would become significant. Previous studies either did not look at effects of gender (Tognin et al., 2012) or found no effect of gender (Klomp et al., 2012). Determining whether the mammillary bodies in men are more vulnerable in schizophrenia, or whether the numerical differences found in the present study might be explained by differences in additional lifestyle factors, is an important future goal.

The current findings differ from previous MRI studies that either reported no difference in mammillary body volume (Klomp et al., 2012) or an increase, driven by changes in the right hemisphere (Tognin et al., 2012). These differences may be due to the boundaries used for mammillary body segmentation and the extent to which adjacent hypothalamic nuclei and/or white matter were included in the measurements. It also could be due to the present study using all planes, rather than just the coronal plane, for segmentation, as well as re-sampling images, which may improve accuracy. The smaller sample size (26 patients versus 26 controls) in the Tognin et al. study (2012) could also contribute to the differing results given variation in mammillary body volumes, even in control populations. In the current dataset there was also variability in the patient group with some patients showing little or no mammillary body atrophy and others showing very marked volume loss. This is consistent with schizophrenia being highly heterogenous and likely encompassing different subcategories, some of which may be more predisposed to mammillary body pathology. Furthermore, there appears to be a complex relationship with age which is likely driven by disease stage and could explain some of the previous contrasting findings depending on the age of participants. In younger patients there was a sharp rise in mammillary body volume which would be consistent which initial inflammation and swelling during early disease stages and the reported volume increase in animal, post-mortem and MRI studies (Briess et al., 1998; Smedler et al., 2022; Tognin et al., 2012) and this is followed by subsequent atrophy during later disease stages. Non-linear changes have previously been reported in longitudinal studies with more dynamic changes, including volumetric increases at the earliest stages of illness (Cropley and Pantelis, 2014). Large-scale longitudinal studies are needed to better understand these variations across patients and age and how they relate to disease progression and symptomology. Furthermore, it would be important to determine whether early markers of mammillary body pathology could be used to better stratify patients for subsequent treatment and neuroprotective protocols and whether there are potential changes within the mammillary bodies prior to symptom onset.

The mammillary bodies receive a dense input from the hippocampal formation (McNaughton and Vann, 2022); given the hippocampus has been repeatedly implicated in schizophrenia, one possibility is that mammillary body atrophy reflects the loss of hippocampal inputs. We obtained automatic estimates of hippocampal and subiculum volumes and found a modest reduction in the hippocampal volumes but no changes for the subiculum. The hippocampal volume changes were consistent with the unadjusted volumes previously reported for this same data set [left and right hippocampus reduced in patients by 4% and 2%, respectively (Zheng et al., 2019)]. This is less than the approximate 10.3% reductions in raw volume seen for the mammillary bodies. The hippocampus was automatically segmented so the differences in degree of volume reduction could reflect differences in accuracy between manual and automatic segmentation although hippocampal segmentation with FreeSurfer has been found to correlate well with manual segmentation in both healthy controls and patients with schizophrenia (Arnold et al., 2014). Alternatively, it may be that the mammillary bodies are more sensitive, and show greater atrophy than the hippocampus, at least in some individuals. Consistent with this, the anterior thalamic nuclei were reportedly affected in first-episode patients, at a point where no differences were observed in the hippocampus (Hoang et al., 2021). The mammillary bodies were also found to be significantly smaller in a genetic rodent model of schizophrenia, with mammillary volume correlating to psychotic symptomology, and this was found to be independent of hippocampal input (Küçükerden et al., 2022).

As there are no background data for the participants in this data set, it is not possible to determine the extent to which any volumetric differences may be exacerbated by lifestyle differences (e.g., increased alcohol consumption), co-occurring anorexia (Meys et al., 2022; Morylowska-Topolska et al., 2017), and/or treatment protocols. However, the changes in mammillary body volume in younger patients, with both atrophy and swelling present across patients, suggests that while the mammillary bodies may show increased vulnerability to lifestyle and/or medication factors, mammillary body pathology is still likely to be a primary feature of schizophrenia. Consistent with this, the anterior thalamus, a brain region closely connected to the mammillary bodies, is also smaller in first-episode psychosis (Hoang et al., 2021). Furthermore, the mammillary bodies show abnormalities in animal models of schizophrenia, highlighting their vulnerability even without concomitant treatment and/or lifestyle differences (Küçükerden et al., 2022; Smedler et al., 2022).

The lack of cognitive data included in current dataset means it is not possible to associate mammillary body volume with cognitive performance. However, previous studies have shown recollective memory performance significantly correlates with mammillary body volume, with poorer memory scores found in patients with smaller mammillary volumes (Annink et al., 2021; Tsivilis et al., 2008; Vann et al., 2009). Other studies have reported correlations between mammillary body volume and anxiety in patients with schizophrenia (Tognin et al., 2012).

Together, the findings from the present data provide further support for the involvement of the mammillary bodies in schizophrenia. The mammillary bodies are associated with both memory and anxiety (Vann, 2010) and mammillary body pathology in schizophrenia could therefore contribute to both the cognitive impairments and changes in affect observed in this condition. A better understanding of the circumstances in which the mammillary bodies are most vulnerable, whether this is a genetic predisposition, treatment protocols, co-morbidities such as anorexia and or/lifestyle differences, would help identify ways to protect the mammillary bodies in schizophrenia and potentially help reduce accompanying symptomology.

## Acknowledgements

This work was supported by a Wellcome Trust Senior Research Fellowship awarded to SDV (WT212273/Z/18). The imaging data and phenotypic information was collected and shared by the Mind Research Network and the University of New Mexico funded by a National Institute of Health Center of Biomedical Research Excellence (COBRE) grant 1P20RR021938-01A2. For the purpose of open access, the author has applied a CC BY public copyright license to any Author Accepted Manuscript version arising from this submission.

